# Application of Pedimap — a pedigree visualization tool — to facilitate the decisioning of rice breeding in Sri Lanka

**DOI:** 10.1101/2020.01.20.912477

**Authors:** P.G.R.G. Rathnayake, S.M. Sahibdeen, U.A.K.S. Udawela, C.K. Weebadde, W.M.W. Weerakoon, S.D.S.S. Sooriyapathirana

## Abstract

The development of rice cultivars with desirable traits is essential. The decision-making is a crucial step in rice breeding programs. The breeders can make efficient and pragmatic decisions if an organized pedigree visualization platform is available for the material of the rice breeding germplasm. The staple food in Sri Lanka is rice, and there is a great demand for improved varieties with high yield and other promising traits. In the present study, the available data of all the rice varieties released by Rice Research and Development Institute, Sri Lanka, and the related landraces and genotypes were arranged in Pedimap, a pedigree visualization tool. Pedimap can showcase pedigree relationships, phenotypic, and molecular data. The Identity by Descent (IBD) probabilities were calculated using FlexQTL software and included in the Pedimap database. The parentage selection based on the variations of phenotypic traits, selection of marker alleles for molecular breeding, and detection of the founders of genetic effects can be swiftly conducted using Pedimap. However, the power of harnessing the value of Pedimap for making breeding decisions relies on the availability of data for the traits, markers, and genomic sequences. Thus, it is imperative to characterize the breeding germplasms using standard phenomic and genomic characterization procedures before organized into Pedimap. Thereby, the worldwide breeding programs can benefit from each other to produce improved varieties to meet global challenges.

## INTRODUCTION

Rice is one of the major crops in the world, with an annual production over 700 million metric tons [1]. Half of the world population consumes rice as the staple food [2]. Currently, the demand for rice is rapidly increasing due to the growth of the human population [3]. However, the current rice production cannot meet the increasing demand causing severe food security issues. The biotic and abiotic stresses also exert a negative influence on rice production [4]. The rice farming is also a way of living for many people, especially in numerous Asian countries [5]. At present, 1.8 million Sri Lankan families engage in rice farming over 870,000 hectares [6]. The annual rice production in Sri Lanka is approximately 2.3 million metric tons (MT), which is insufficient to fulfill the domestic rice demand of 3.0 million [7]. Hence, the Sri Lankan government spends about USD 400 million to import rice annually [7,8].

The rice production is mainly affected by drought and irregular rainfall patterns caused by climate change [9,10,11], adverse soil conditions such as salinity [6,12], and pest and disease attacks [13]. The biotic and abiotic stresses in rice farming can be controlled using numerous agronomic practices such as irrigation, drainage, fertilization, and the application of pesticides. However, the rate of success of the controlling methods is limited [13] due to the unpredictable nature of climate change, soil degradation, variations in pest dynamics, and development of pest resistance [14]. Therefore, breeding is considered as the most successful strategy to produce high yielding and stress resilient rice varieties [15]. The improved rice genotypes can also contain the traits for higher consumer preference and organic farming [16]. In the past, the rice varietal improvement was conducted with classical breeding techniques, which are tedious, lengthy, and less feasible in cases such as breeding for pest resistance and submergence tolerance. However, the marker-assisted breeding (MAB) is employed in modern breeding programs to introgress valuable genetic loci from landraces and traditional varieties [17,18] and the desirable haplotypes of Quantitative Trait Loci (QTL) to the improved rice varieties [19–21].

The decision-making process in a breeding program is crucial for successful outcomes. The formulation of decisions before breeding is a multi-step process that consists of the identification of breeding priorities, determination of the genetics of target traits, and employment of pre-breeding methods if required. The economic and technical feasibility, number of parents for crosses, number of selfing and outcrossing cycles, length of the breeding program/cycles, and identification of the selection methods must also be assessed [22]. In the decisioning process, initially, the market trends based on consumer and other stakeholder preferences must be recognized [23]. Subsequently, the novelty and the uniqueness of the breeding objective must be assessed before the execution of the breeding program [22].

The selection of suitable varieties or individual plants as parents and the determination of the selection methods are the two most critical aspects in planning breeding programs [24]. The parental selection depends on the number of prioritized traits for breeding. When multiple characteristics are to be introgressed, the breeders require a prioritized order of parents for stepwise crossing and selection [25,26]. The decision-making process in breeding is entirely based on the available information on phenotypes, genotypes, pedigree, available budget, field and greenhouse space, desired time-to-market etc. Although the data for decision-making for breeding are indispensable, haphazardly collected information would provide less value to the breeders. In many conventional breeding programs, most of the data are recorded in field notebooks and stored in the breeding stations, while very little information is available as computerized databases. If an organized database containing all the essential information for the rice varieties released and the parental genotypes used in breeding, the decisions can be easily made.

The construction of a database with all the necessary information from varieties and their parents promotes the capacity of data sharing, mining, visualization, and retrieval [27]. Pedimap is a pedigree visualization software. The data needed can be imported to Pedimap from FlexQTL, or with some custom script from any other database program. Pedimap is used by many contemporary plant genetics and breeding programs worldwide. As stated in Voorips et al. (2012) [28], Pedimap can be used to record and utilize breeding history. Pedimap illustrates the available phenotypic and genetic data through pedigrees. All the information, including parentage, qualitative and quantitative data, marker alleles/genotypes, and the calculated identity-by-descent (IBD) probabilities can be presented in Pedimap. Currently, breeders prefer to use pedigree visualization tools like Pedimap since it allows them to access the large pool of genetic and phenotypic data quickly and generate pedigrees that are essential in making breeding decisions.

In Sri Lanka, Rice Research and Development Institute (RRDI) is the sole organization conducting the rice breeding programs for the national needs. Therefore, in the present study, we report an attempt to organize the information of the released varieties and the parental genotypes of RRDI breeding programs as a Pedimap based database which is a valuable step to take accurate breeding decisions and speed up the process of releasing novel varieties.

## MATERIALS AND METHODS

### Data Curation

The data were collected from RRDI, Sri Lanka and classified under three main categories, namely pedigree history, phenotypic data, and molecular data on rice varieties/landraces/genotypes (herein after collectively referred to as cultivars). The male and female parents and the order of crosses were taken under pedigree history. The average yield of the rice plants, the maturity period in different growing seasons (*Yala* and *Maha* seasons of Sri Lanka [29]), plant height, basal leaf sheath color and additional color patterns, recommended type of the land, level of phosphorus deficiency tolerance, amount of brown rice recovery, milling recovery, head rice recovery, amylose content, gelatinization temperature, weight of 1000 grains, shape of the grain, pericarp color, weight of a kg, color of the buff coat and resistance/susceptibility to pests and diseases; brown planthopper (BPH), bacterial leaf blight and rice blast disease were recorded under phenotypic data (S1 Table). The available DNA marker alleles, marker positions in the linkage map, and allelic scores were entered under molecular data [30–33] (S2 Table).

### Pedimap Procedure

A Pedimap input data file is created in MS Excel (2019), and the data file is exported as a tab-delimited text (.txt) file (S3 Table). The input file contains four main subdivisions; header, pedigree, marker data, and IBD probability section (Fig 1). The header consists of five essential elements and one additional element. The name of the population and symbols for unavailable or missing data, null homozygous alleles, and confirmed null alleles are entered to the pedigree section, as shown in Fig 1A. The name of the cultivar must be a string with text or numerical values without spaces.

**Fig 1.**
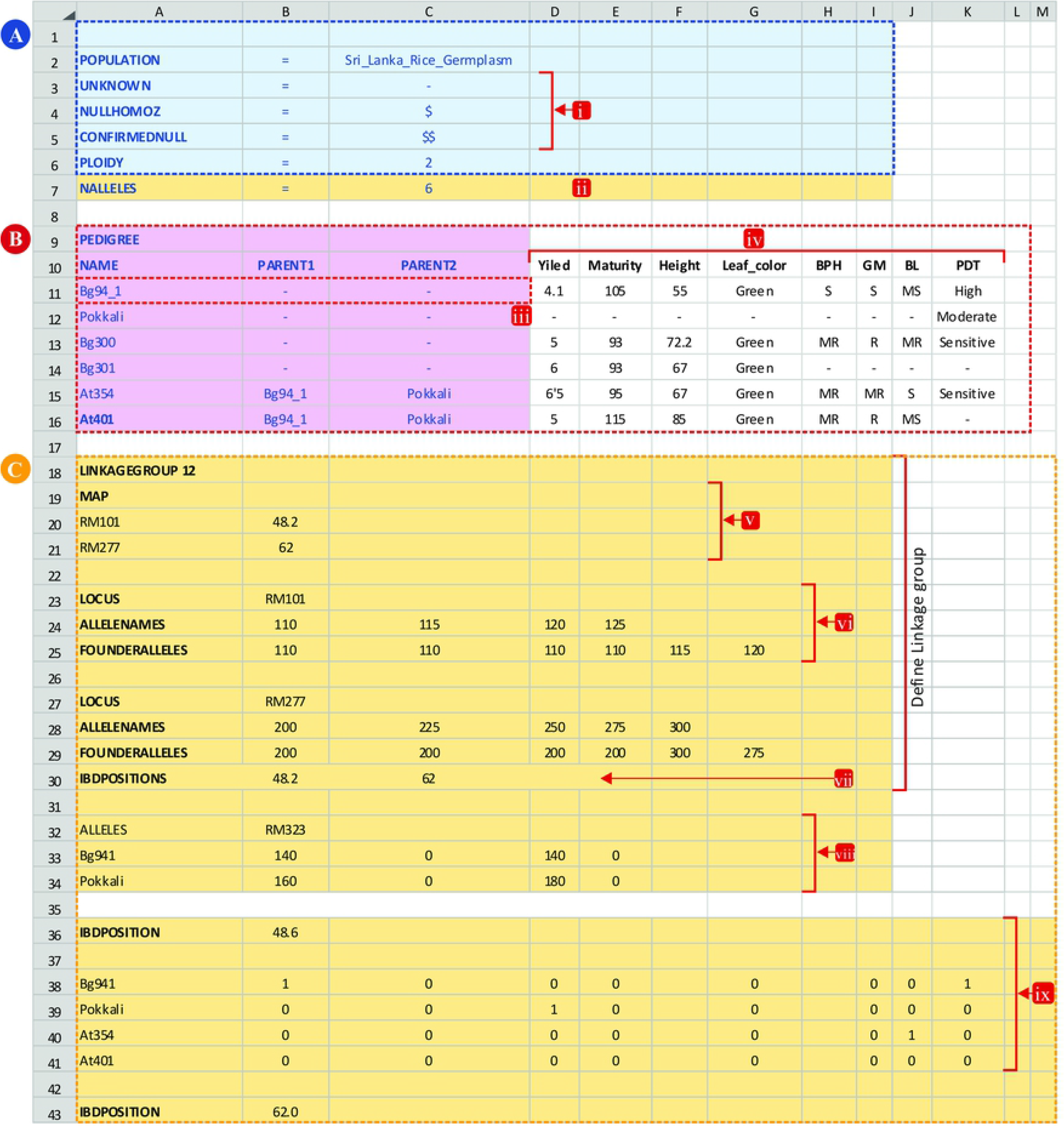
The input data file structure of the Pedimap; The input file was created as an MS Excel worksheet, contains four main sections. A: Header, B: Pedigree and phenotypic data, C: Genotypic data. A: In the header section, essential elements are highlighted in blue, which contains the population name, ploidy and codes used in the data. (i): abbreviations for missing data (i.e., unknown), possible null alleles, confirmed null alleles; (ii): NALLELES is only necessary if the IBD probabilities are used, and specifies the total number of founder alleles (i.e. the number of founder times the ploidy). B: The Pedigree section contains the pedigree data of all the individuals, and any phenotypic data of the individuals. The pedigree part is highlighted in purple. (iii): founders (initial parents) are entered with missing values for their parents. Phenotypic data are entered in subsequent columns (iv). C: The Genotypic data section (if present) is divided into three parts: one part for each linkage group the genetic map (v), general information per locus (vi) and positions where IBD probabilities are calculated (vii); a part with the observed alleles per locus per individual (viii), and a part with the Identity-by Descent (IBD) probabilities per position per individual (ix). The final file must be saved as a text (.txt) file.

Next to the header, the pedigree section is entered, as shown in Fig 1B. The first column denotes the name of the variety or landrace, and second and third columns are reserved for maternity and paternity information, respectively. The numbers and strings can be included to represent the phenotypes in the first three columns. From the fourth column onwards, any desirable quantitative or qualitative trait values can be entered. All the collected phenotypic data are introduced, as shown in Fig 1B. The third section of the input data file is for marker information. The linkage group of the DNA marker and the marker positions in the linkage map are entered, as shown in Fig 1C. If there are more than one linkage group, all the linkage group maps should be defined successively before entering the allelic scores. The detailed data for each DNA marker can be inserted after revealing the map positions. The respective number of columns, according to the ploidy level, should be incorporated to enter allelic scores. The fourth section is for IBD probability values (Fig 1C). The IBD probabilities cannot be calculated within Pedimap but can be calculated using other software *e.g*. FlexQTL [34], which is a software for QTL analysis. FlexQTL can also generate a complete Pedimap input data file.

### Demonstration of the Usability of Pedimap

We used the examples 1 and 2 given in Table 1 to show how parental cultivars can be selected for crossing based on diverse breeding objectives and the prioritized traits. The example 3 in Table 1 was used to select parents, indicate the DNA marker allelic representation for MAB, identity by descent calculations, and planning crosses to deduce related details necessary for decision-making for breeding.

**Table 1.** The examples used to demonstrate the use of Pedimap in making breeding decisions.

| Example | Trait* |  |  |  |  |  |
| --- | --- | --- | --- | --- | --- | --- |
|  | Priority# 1 | Priority 2 | Priority 3 | Priority 4 | Priority 5 | Priority 6 |
| 1 | White pericarp | Yield $\geq 3.5$ mt/ha | Resistance or moderate resistance to brown planthopper (BPH) | Maturity period $\leq 125$ days | Grain shape<sup>§</sup> | - |
| 2 | High, high-intermediate, and intermediate amylose content | Yield $\geq 3.5$ mt/ha | Maturity period $\leq 125$ days | Resistance or moderate resistance to blast | - | - |
| 3 | Phosphorous deficiency tolerance | Yield $\geq 5.0$ mt/ha | Maturity period 90-105 days | Resistance or moderate resistance to BPH | Resistance or moderate resistance to blast | High, high-intermediate, and intermediate amylose content |

\*The trait classes and records are from RRDI records [35].
#The traits are given in the order of priority in making breeding decisions.
§Variable grain shapes of the intended varieties to be released.

## RESULTS AND DISCUSSION

Worldwide plant genetics and breeding programs use Pedimap as the platform for maintaining breeding databases and pedigree visualization. In the RosBREED project [36], the parental and progeny identification, tracing founders, and calculation of allelic representation are conducted using Pedimap. The pedigree display of Pedimap is used to plan crosses in the Rosaceae research community [37, 38], HIDRAS project [39] and visualize of *Arabidopsis thaliana* crosses [40]. Selecting parentage, sketching out crossing schemes, estimating the probability of allelic segregation, and choosing compatible molecular markers for MAB can be achieved using Pedimap [28]. The use of Pedimap as a pedigree visualization tool for the decision-making process in rice breeding is described using three examples (Table 1).

### Example 1: Selecting parents for higher yield, BPH tolerance, short duration and white pericarp with diverse grain shapes

The Pedimap database rice breeding gerplasm in has a total of 224 input cultivars. There are 36 intermediate genotypes such as F1 and F2 that were not reported, but we included them to complete the pedigree in Pedimap. Thus, the database has a total of 188 rice cultivars and accessions with known identities with records (S1 Fig, S1 Table). In Example 1, we considered a scheme to select accessions as parents with the parameters given in Table 1 for white pericarp, yield, BPH resistance, maturity period, and the grain shape. These thresholds defined a subpopulation of 26 cultivars (Fig 2). The variation of the yield is given in Fig 2A. According to the color shading given, the breeder can select the required parents for crossing to obtain higher yield levels. However, as shown in Fig 2B, only three cultivars show the complete resistance to BPH. If breeder plans to introgress the complete BPH resistance to the novel varieties, only Bg250, At307, and At306 are available as the sources of resistance. Fig 2C displays the variation for the maturity period. The breeder can choose the parents depending on his objective for the intended maturity period for the novel varieties. Example 1 was exclusively planned to breed for white pericarp. However, the grain shape is also important as a significant quality trait to become a successful variety in the market. Fig 2D shows the variation for grain shapes for the breeder to carry out the selection. If we consider all the traits and selected At307 as a parent based on the pedigree visualization in Pedimap, At307 can provide the genetic basis for high yield, complete resistance to BPH, approximately three months for maturity and intermediate-bold shaped grains. If Bg450 was selected, the yield is still in the higher range with moderate resistance for BPH and short-round grains. However, Bg450 brings the alleles for an extended maturity period (Fig 2).

**Fig 2.**
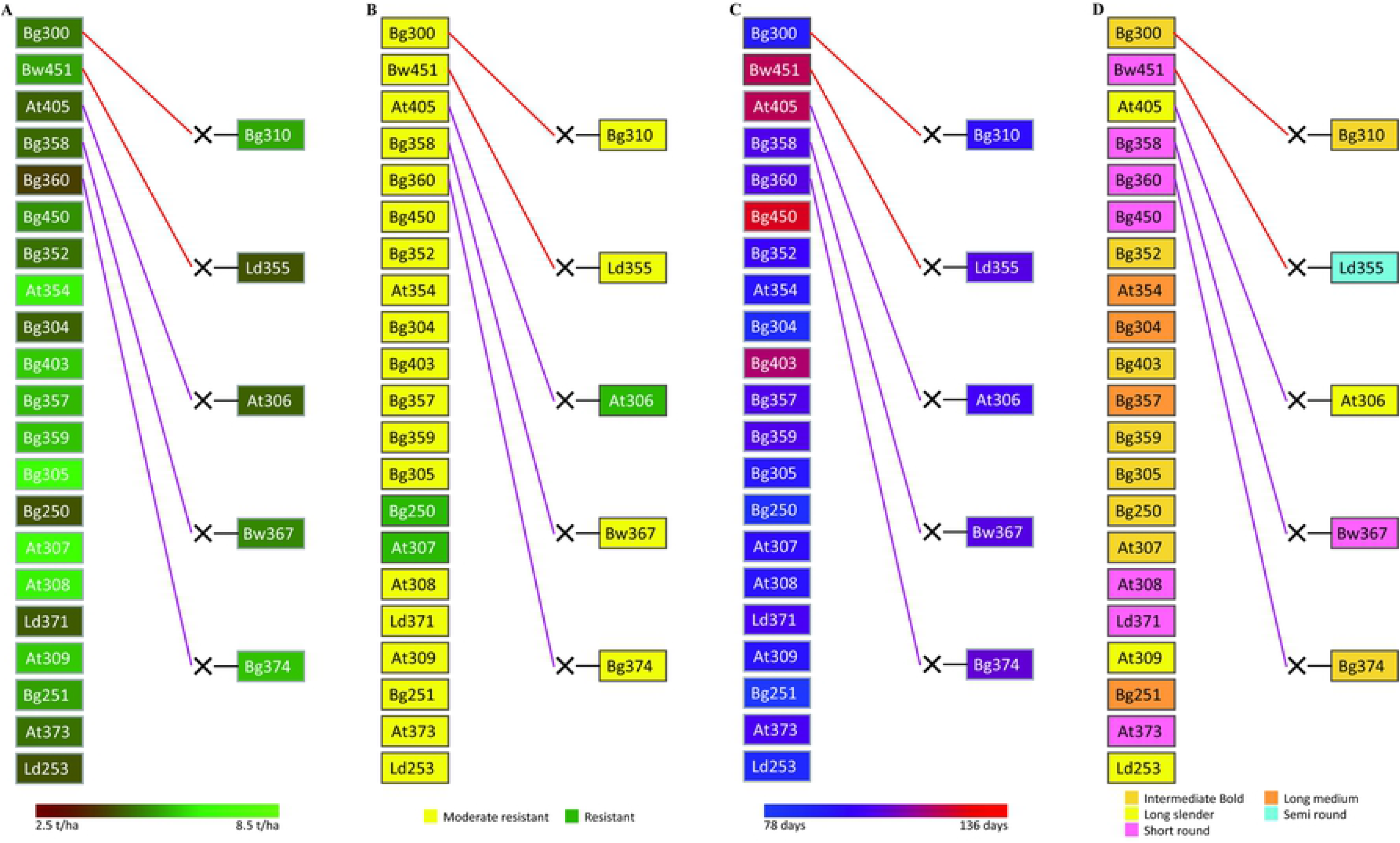
The pedigree visualization for Example **1 (**Parents with white pericarp, yield ≥ **3.5** mt/ha, moderate or complete BPH resistance, maturity period ≤ **125** days, and diverse grain shapes). The selected pedigree is colored separately for four traits. A: Yield; B: Degree of resistance to brown planthopper (BPH); C: Maturity period; D: Grain shape. Female and male parentages are indicated by red and purple lines, respectively. The symbol ‘×’ indicates the cross between two parents. The background colors of the cultivar-name boxes indicate the trait values, as shown in the colored legends below.

### Example 2: Selecting parents for high/high-intermediate amylose content, higher yield, short duration, and resistance to blast disease

In Example 2, we considered a scheme to select cultivars/accessions as parents with the parameters given in Table 1 for high/high-intermediate amylose content, higher yield, short duration, and resistance to blast disease. These thresholds defined a subpopulation of 37 cultivars/accessions (Fig 3). The breeder can select the high yielding, short-duration, and blast-resistant cultivars as parents from pedigrees visualized in Figs 3A, 3B, and 3C, respectively. The high, high-intermediate, and intermediate amylose contents are depicted in the pedigree given in Fig 3D. Only Bw351, At307, Bg407H, At308, and Bg252 show the complete resistance to blast (Fig 3C). However, At307 is the most promising parent with high yield (Fig 3A), short duration (Fig 3B), and high amylose content (Fig 3D) along with complete resistance to blast (Fig 3C). Also, Bg407H is the highest yielding (Fig 3A), blast-resistant (Fig 3C), and high in amylose content (Fig 3D). However, Bg407H is a long duration variety compared to At307. Therefore, the breeder may plan to cross At307 and Bg407H to accomplish the breeding objective of Example 2.

**Fig 3.**
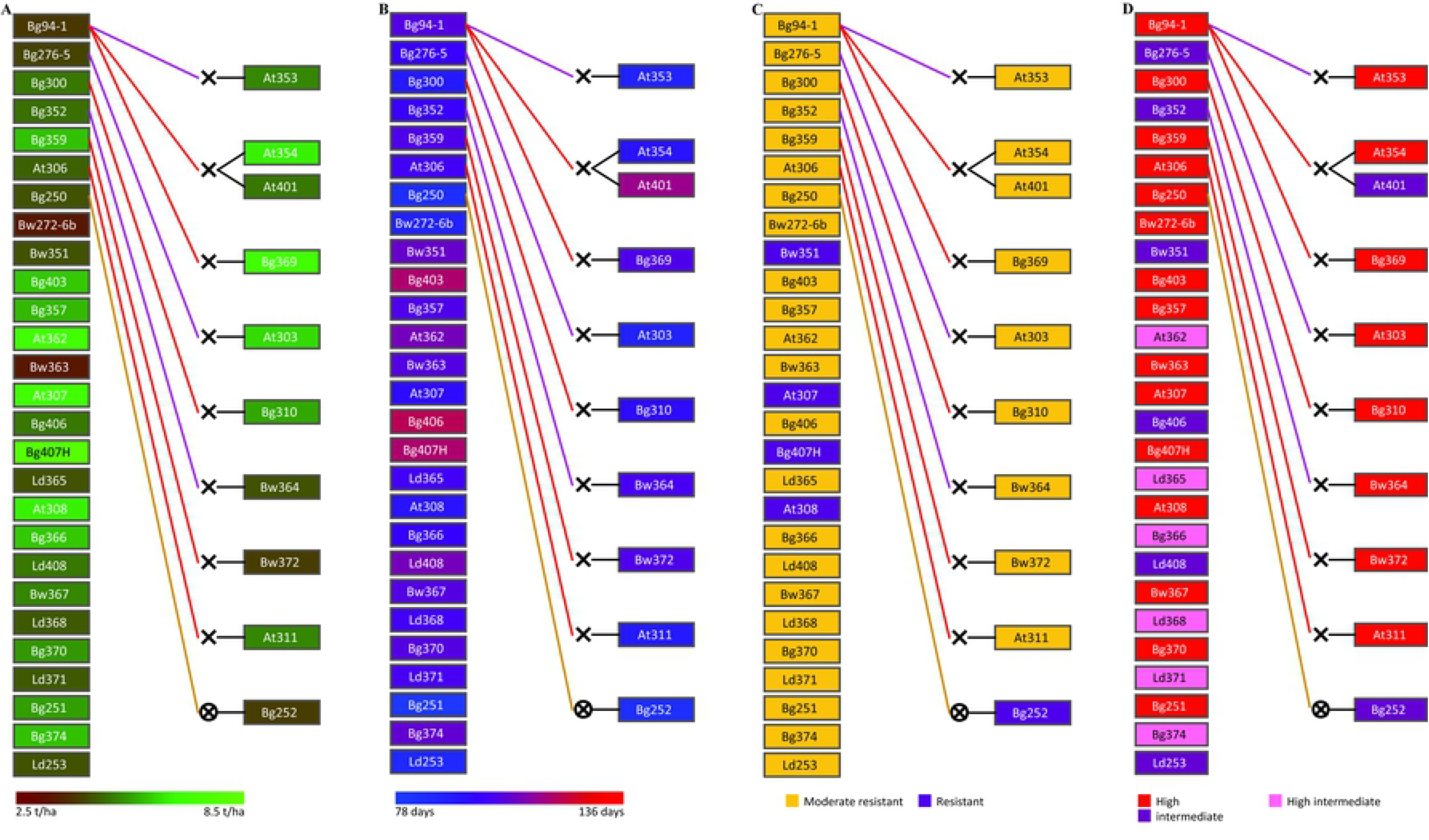
The pedigree visualization for Example 2 (parents with high, high-intermediate, and intermediate amylose content, yield ≥ 3.5 mt/ha, moderate or complete resistance to rice blast disease and maturity period ≤ 125 days). The selected pedigree is colored separately for four traits. A: Yield; B: Maturity period; C: Degree of resistance to rice blast disease; D: Amylose content. Female and male parentages are indicated by red and purple lines, respectively. The symbol ‘×’ indicates the cross between two parents, and ‘×’ inside the circle represents selfing. The background colors of the cultivar-name boxes indicate the trait values, as shown in the colored legends below.

### Example 3: Selecting parents for phosphorus deficiency tolerance, higher yield, short duration, resistance to both BPH and blast, and high/intermediate-high amylose content

We selected a set of rice cultivars from the Pedimap database based on the availability of ranked scores for phosphorus deficiency tolerance (PDT). Twenty-four cultivars contain the PDT ranks of high, moderate, and sensitive (Fig 4A). The same set was illustrated using Pedimap for yield (Fig 4B), maturity period (Fig 4C), degree of resistance to BPH (Fig 4D) and blast (Fig 4E), and amylose content (Fig 4F). If At362 is considered as a parent, it can bring resistance to PD, and BPH, moderate resistance to blast, high yield, average maturity period, and intermediate-high amylose content. Similarly, if Bg250 is selected, it can bring moderate resistance to PD and blast, resistance to BPH, moderate yield and shortest maturity period, and high amylose content (Fig 4).

**Fig 4.**
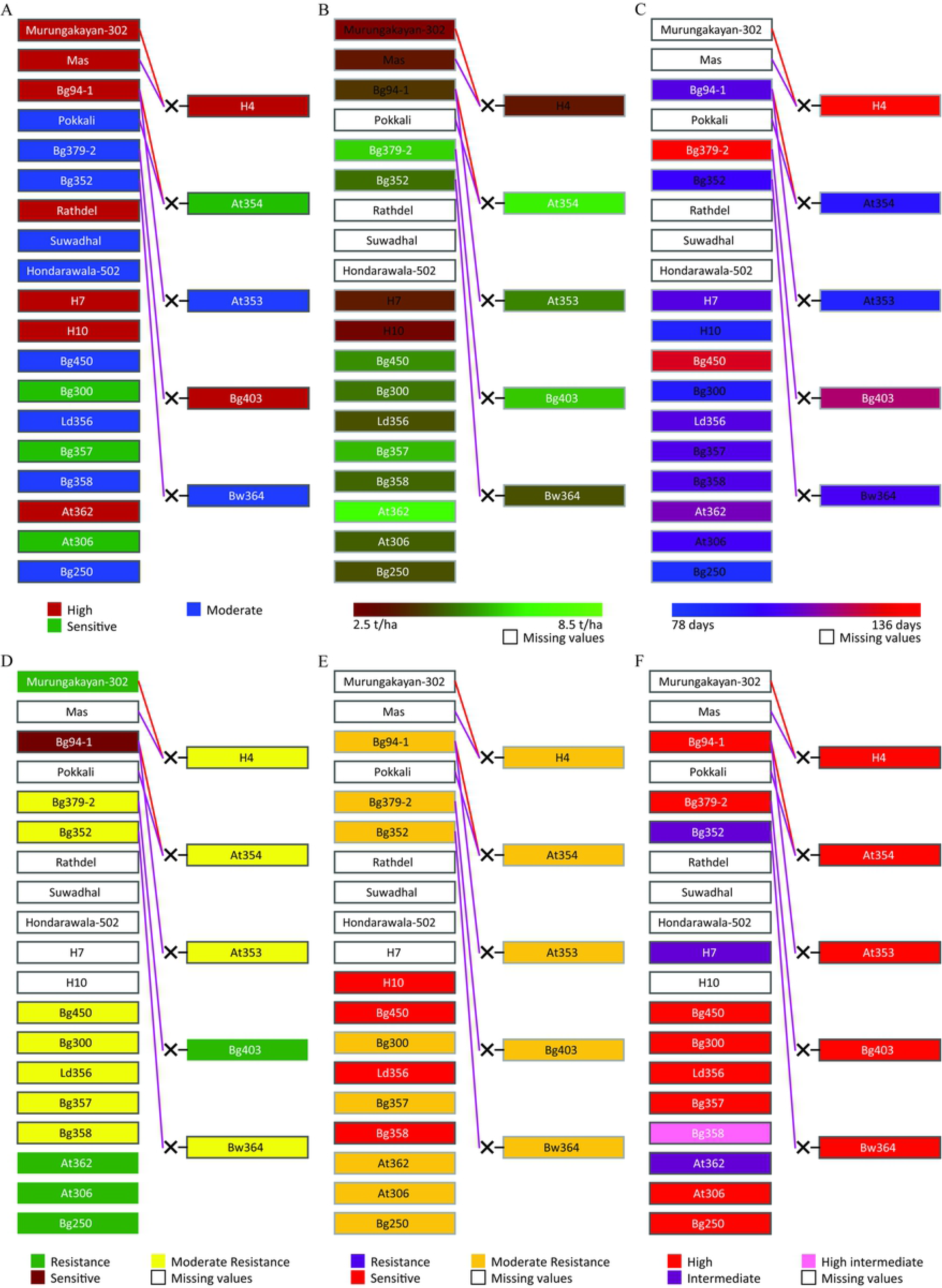
The pedigree visualization for Example 3 (parents ranked for phosphorus deficiency tolerance). The selected pedigree is colored separately for six traits. A: PDT; B: Yield; C: Maturity period; D: Degree of resistance to BPH; E: Degree of resistance to BLAST; F: Amylose content. Female and male parentages are indicated by red and purple lines, respectively. The symbol ‘×’ indicates the cross between two parents. The background colors of the cultivar-name boxes indicate the trait values, as shown in the colored legends below. The cultivars with missing-trait values are indicated by white boxes.

A sample crossing scheme is shown in Fig 5 to produce a rice variety with high PDT, mean yield ≥5.0 mt/ha, maturity period ≤105 days, resistant to BPH and blast disease, and higher amylose content. Since there is no reported cultivar for high PDT with complete blast resistance (Fig 4), the illustrated crossing scheme in Fig 5 is proposed with two phases. In the first phase, the crossing of At362 and Bg250 followed by numerous rounds of selfing and selection of the most beneficial lines among the RILs at advanced generations would accomplish the breeding objective only without complete resistance to blast (i.e., a moderate level of blast resistance is possible). In the second phase, the selected RILs from phase 1 can be backcrossed to Bg252 as the donor parent to introgress the complete resistance to blast. The breeder can come up with diverse crossing schemes like the one given in Fig 5 to make effective decisions for breeding and maximize the resource utilization to release varieties in the shortest possible time. The breeder can select any number of parents that are needed to use as sources of resistance and other traits to start crossing. Also, the marker alleles and the IBD probabilities can be checked as illustrated in S2A Fig and S2B Fig respectively.

**Fig 5.**
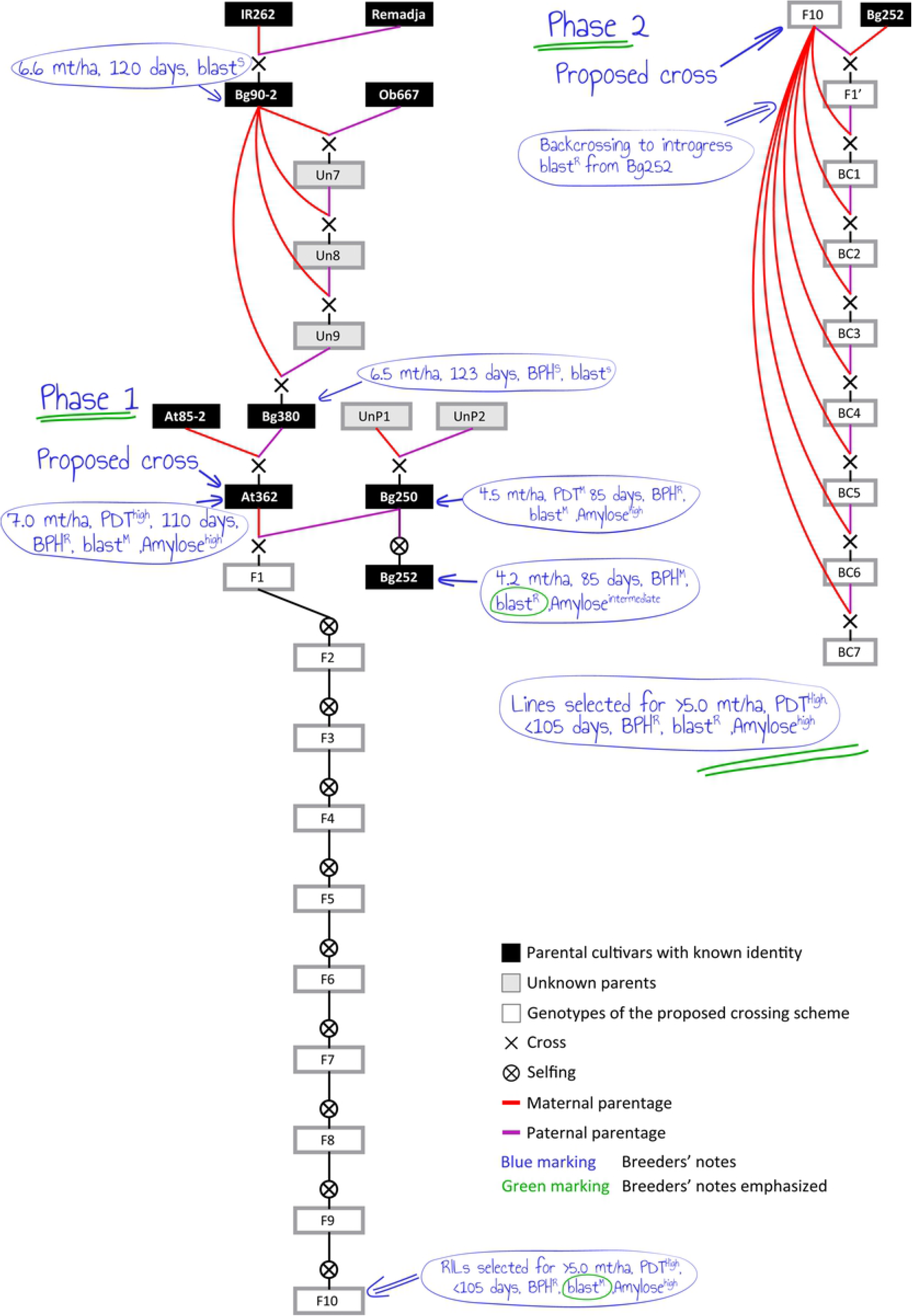
The pedigree visualization for planning a crossing scheme. Phase 1: Initial crossing of At362 and Bg250 and pedigree selection to obtain RILs with ≥5.0 mt/ha of mean yield, ≤105 day of maturity period, resistant to BPH, moderately resistant to blast and high level of amylose content. Phase 2: Then backcrossing with Bg252 as the donor parent to introgress the blast resistance.

The decision-making process in breeding is a tedious task [22]. The breeding germplasm is complex with large numbers of improved varieties, traditional cultivars, landraces, wild germplasm and accessions. Also, there can be large mapping populations and unreleased varieties due to various reasons. The numerous genotypes in breeding germplasm may have extensive records on agronomic data, pest and disease resistance, quality traits, availability of samples, geographic locations, and utilization in diverse breeding programs as parents [41, 42]. With the advent of DNA markers and sequencing technologies, a wealth of genomic information is also available [43]. However, one of the recurrent problems in any breeding germplasm in the world is most of the cultivars remain uncharacterized. Thus, they cannot be used directly in breeding activities. Traditionally, breeders keep records in field books. With the development of computer technology, data tabulation is becoming a common practice. However, given the highly complex nature of the datasets in breeding germplasm, data tables have a limited value to the breeders. The tables created with contemporary data managing software cannot graphically display complex pedigrees and variations of qualitative and quantitative traits along with DNA marker information. These database handling platforms do not make use of the pedigree-based capabilities of Pedimap, like selecting related parental varieties/accessions. In this context, Pedimap provides a considerable advantage, as it can visualize pedigree relationships, trait variations, and any other useful information required for decision-making and planning crosses in breeding programs [28]. If all the available details on breeding germplasm are arranged as a database, the breeder can come up with subpopulations based on diverse traits and select the parents for improving multiple traits. However, simple spreadsheets or manually prepared note pages cannot be used to visualize the essential information and complex pedigrees. Breeding programs often suffer a lot when the breeder gets retired or moved to a different position [44–46]. The newly hired breeder cannot practically go through the individual records of the existing breeding germplasm. Thus, there is a strong possibility that valuable breeding germplasm might get lost wasting time, resources, and courage of the retired breeder and his team. However, as a routine practice, if the breeder maintains and updates a Pedimap file for the developing germplasm of breeding materials, the newly hired breeders can go through and identify the value and gaps in the available material for him to plan further. The creation of a Pedimap file is simple, and a novice to informatics can curate and use Pedimap with a little training. Pedimap allows breeders to store data, fetch and visualize genomic information at any time with less effort and complete accuracy [47]. The straightforward accessibility, direct data interpretation, ability to customize the views in multiple fashions, and editable output file formats are the significant features of Pedimap. The graphic files created can be readily imported to image editing software for further visualizations and illustrations. Pedimap is not an opensource software but can be freely obtained by contacting the developers thus even the breeders in developing countries can benefit from Pedimap [28].

In the current study, we created a Pedimap database for the rice cultivars and accessions prominently used by breeding programs in Sri Lanka. With the available information, significant breeding decisions can be made as we explain in three examples (Figs 2–5). However, it is essential to characterize the cultivars for all the important traits, molecular markers and SNP haplotypes [48], so that breeding decisions can be effectively made [17]. The phenotyping methods must be standard and should follow common procedures across different locations so that the power of the Pedimap database would go up dramatically. Therefore, breeders should always follow the standard, globally acceptable phenomic platforms to characterize the material in breeding germplasm [39, 49].

## CONCLUSION

The pedigree visualization with variations of phenotypic and molecular data using Pedimap is a user-friendly tool to plan rice breeding programs with higher accuracy and resource optimization. The present study explains the applicability of Pedimap as a decision-making tool to streamline the rice breeding programs in Sri Lanka. However, it is also important to note that accurate characterization of the breeding germplasm for phenotypic and molecular data is the critical prior step to harness the value of Pedimap for breeding.

## Supporting information

**S1 Table** Varietal data

**S2 Table** Marker data

**S3 Table** Pedimap input data file

**S1 Fig** Visualization of the entire pedigree of the rice cultivars in the rice breeding germplasm of Sri Lanka. Female and male parentages are indicated by red and purple lines, respectively. The symbol ‘×’ indicates the cross between two parents, and ‘×’ inside the circle represents selfing.

**S2 Fig** Visualization of selected marker genotypes and Identity by Descent (IBD). A: Marker alleles. The alleles of the DNA markers K29-N, K41, K48, and K5-N are given in vertical order.; B: IBD probabilities of four Pup1 linked markers. The same pedigree as in the S2A Fig is shown with precalculated IBD data for part of the Pup1 linked region on chromosome 12 at about 55 cM. Since the cultivar linkage maps are not available, we assumed a 0.1 cM gap between adjacent markers for the representation of IBD values. Eachcolor represents a different founder, haplotype. Each rectangle represents one copy of the selected chromosome in an individual. The chromosomal position of the alleles represents the vertical bars, and the width of a color bar indicates the IBD probability of the corresponding founder alleles.

